# VRPG: an interactive visualization framework for reference-projected pangenome graph

**DOI:** 10.1101/2023.01.20.524991

**Authors:** Zepu Miao, Jia-Xing Yue

**Author notes:** Corresponding author: (J.-X. Y.). E-mail: Zepu Miao >, Jia-Xing Yue >.

## Abstract

With the increasing availability of high-quality genome assemblies, pangenome graph is emerging as a new paradigm in the genomics field for identifying, encoding, and interpreting genomic variation landscape at both population and species levels. However, it remains a major challenge in understanding and interpretating pangenome graph via biologically informative visualization. To facilitate a better examination of pangenome graph towards novel biological insights, here we present VRPG, a web-based interactive visualization framework of pangenome graph. VRPG provides dynamic and efficient supports for exploring and annotating pangenome graphs along a reference-based coordinate system. Moreover, VRPG shines with its unique features in highlighting the directed path of any graph-constitutive assembly as well as in denoting copy number variation of genomic segments represented by graph-constitutive nodes, which are highly valuable in genome comparison and variant analysis. Additionally, VRPG enables side-by-side visualization between pangenome graph and reference-based annotation features, bridging the graph and linear genome context. To further demonstrate its features and scalability, we applied VRPG to the cutting-edge yeast and human reference pangenome graphs derived from hundreds of high-quality genome assemblies via a dedicated website (https://www.evomicslab.org/app/vrpg/).

**Key points:** Developed an intuitive visualization framework for pangenome graph.

Empowered graph-based feature highlighting and variant visualization.

Enabled connecting reference-based annotation features in the context of pangenome graph.

## Introduction

Long-read-based DNA sequencing has become the go-to-choice for most genome sequencing projects nowadays, empowering the production of chromosome-level telomere-to-telomere (T2T) genome assemblies for diverse organisms (including human) [1–3]. With such T2T reference assembly panel continuously expanding at both population and species levels, researchers began to use pangenome graphs to better represent the population- and species-wide genomic variation landscapes in a sequence-resolved manner [4]. Compared with the conventional linear reference genome, pangenome graph offers enhanced power and accuracy in read mapping and variant discovery, especially in the presence of sequence polymorphisms and structural variants (SVs) [5]. Therefore, a representative pangenome graph is expected to shed novel insights into the interpretation of the genotype-to-phenotype association and the discovery of missing heritability.

While a number of tools have been developed to build pangenome graph based on genome alignments, Minigraph [6], Minigraph-CACTUS [7], and PGGB [8] are among the most popular ones. Minigraph is designed for efficiently constructing a reference-based pangenome graph with large variants (e.g., SVs) compactly encoded in its accompanied reference Graphical Fragment Assembly (rGFA) format, which records the origin of each graph constitutive segment from the input linear genomes and therefore allows for traversing the pangenome graph along a stable coordinate system. This unique feature makes the pangenome graph built by Minigraph a natural and intuitive extension to the conventional linear reference genomes, although certain level of bias might be introduced during the graph building process regarding selected reference and input genome order [9]. Minigraph preferentially considers large variants (e.g., SVs) while being less discriminative for small variants such as single-nucleotide-variants (SNVs) and small insertion/deletions (INDELs) during graph construction, therefore pangenome graphs created by Minigraph is not a strictly lossless representation of the full genetic variation possessed by the input genomes at the per base level. As an improvement, Minigraph-CACTUS was proposed as a new pangenome graph building pipeline that combines the compactness and efficiency of Minigraph as well as the base-level sensitivity and accuracy of CACTUS. Therefore, the pangenome graph built by Minigraph-CACTUS can effectively represents the full genetic variation across both base-level and structural-level scales while still preserving the coordinate of the linear reference genome. Finally, PGGB adopted a reference-free approach for pangenome graph construction based on all-against-all genome alignments, which is theoretically unbiased but computationally more expensive. Also, PGGB will clip and collapse the highly similar genomic segments of the input genomes and therefore disrupt the linearity of genome coordinates, making it less straightforward in tracing the original genome coordinates of each graph constitutive segment.

An intuitive visualization of pangenome graph can greatly assist researchers to explore and understand the global and local genomic variation in their graph representation. To date, several tools have been developed for visualizing pangenome graph, among which Bandage [10] and GFAviz [11] focus more on large-scale topology, while SequenceTubeMap [12], MoMI-G [13] , and PGR-TK[14] are more suitable for visualizing fine-scale sequence level details. ODGI [15] showed improved performance on large-scale pangenome graphs with extended visualization function for binned and linearized 1-dimentional local graph structure rendering. However, none of these tools are suitable for dynamically visualizing full-scaled pangenome graphs on the fly due to limitations in scalability. Moreover, most of these tools only provide simple visualization function for the graph itself with limited features in exploiting the connection between pangenome graph and the conventional linear-genome-based annotation features.

In this study, we present VRPG, a web-based interactive visualization framework for reference-projected pangenome graph with scalability. VRPG natively supports the rGFA-formatted pangenome graph built by Minigraph and provides pre-shipped auxiliary command line module (gfa2view) for projecting pangenome graphs built by Minigraph-CACTUS and PGGB (both in GFA format) to a reference-based stable coordinate system. Such design helps VRPG to render a linear reference-based intuitive pangenome graph visualization while providing biological informative features such as genome path highlighting and copy number variation presenting, which are highly valuable in both large-scale genome comparison and fine-scale variant analysis. Moreover, VRPG is algorithmically optimized to be highly scalable, capable of reading, processing, and dynamically rendering large-scale pangenome graphs built upon hundreds of input genome assemblies. Finally, an additional genome annotation track is natively supported with VRPG, which allows for associating the pangenome graph with linear-reference-based annotation features, providing a highly intuitive way of exploring functional genomic features in the context of pangenome graph. As a live demonstration, we applied VRPG to the cutting-edge yeast and human pangenome graphs via a dedicated website (https://www.evomicslab.org/app/vrpg/), which showcases the feature and power of VRPG in exploring the power of pangenome graph.

## Results

### Software design

VRPG is an interactive web visualization framework for pangenome graphs encoded in rGFA or GFAv1 formats. VRPG is easy to deploy and highly efficient in visualizing complex pangenome graphs derived from hundreds of genome assemblies on the fly. VRPG renders pangenome graph by implementing a reference-based projection, which make it highly intuitive and informative to relate the pangenome graph topology to biological annotations compiled for the traditional linear reference genome. Moreover, a graph simplification function is further implemented to enable users to dynamically remove nodes corresponding to small variants (e.g., SNVs and INDELs), which is very useful when visualizing pangenome graphs built by Minigraph-CACTUS and PGGB, in which base-level small variants are encoded as individual nodes and therefore making the graph topology become too complex to display as the query region extends.

VRPG mainly operates via a web browser, in which users can easily query and browse the input pangenome graph (Figure 1). Once users specify the query region based on its genomic coordinates in the reference genome, VRPG can extract and render the corresponding subgraph in three different layouts: squeezed (default), expanded, and cola. Three levels of graph simplification strategies are provided: non-ref nodes (default), all nodes, and none. Users can select the preferred strategy based on the graph complexity of the query genomic region. In the rendered graph, further information regarding the node’s genomic origin, covered gene list, copy number status, as well as the assembly-to-graph path can be easily extracted and displayed by simple clicks, which comes handy for users to relate the graph topology to the input linear genome assemblies. The reference-based annotation file can be visualized together with the pangenome graph in a side-by-side manner.

**Figure 1.**
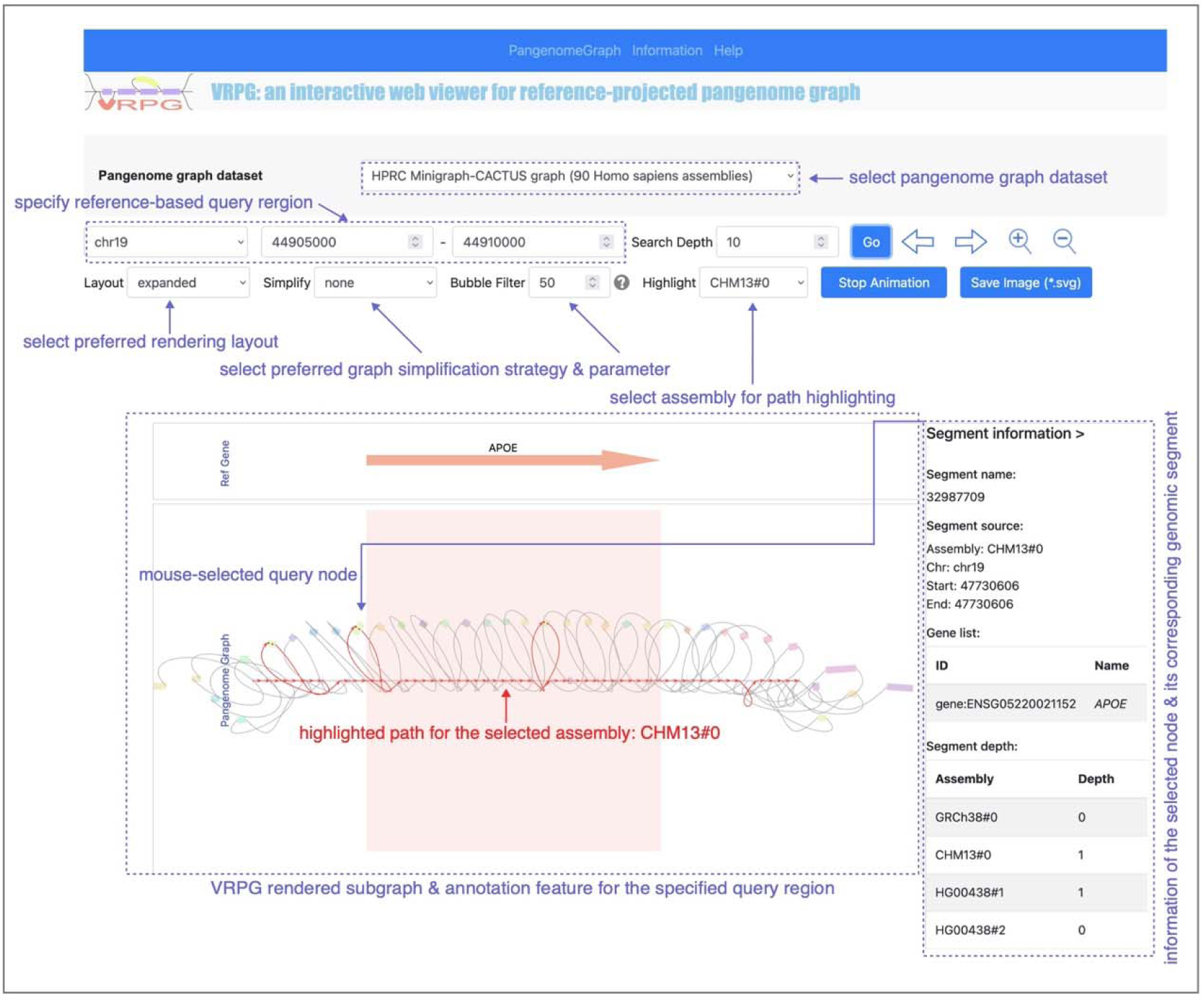
The interactive user interface of VRPG when opened in a web browser. After selecting the input pangenome graph dataset, users can interactively visualize, navigate, and query the pangenome graph based on reference-based genomic coordinate system. The human *APOE* gene region in the HRPC Minigraph-CACTUS pangenome graph is shown here as an example. The *APOE* gene is a clinically important gene bears strong associations with cardiovascular and Alzheimer’s diseases.

### Basic view and highlighting

In the graph rendered by VRPG, graph nodes correspond to the pre-defined reference genome are shown along the center line, while non-reference nodes are shown in the peripheral space. When clicking on a graph node, the node block will become thicker and the information about th corresponding genomic segment will be reported on the right panel (Figure 2A). When clicking on a graph edge, the edge will be colored in red (Figure 2B) and its color will revert to black on a second click. In addition, VRPG provides a “Highlight” feature for highlighting the path of any constitutive genome assembly in the pangenome graph (Figure 2C). When selecting a specific genome assembly in the “Highlight” field, the assembly-to-graph path of the selected assembly will be highlighted in red. All matched nodes will be marked by red dotted lines and all matched edges will be marked by red solid lines. The arrows on the matched edges reflect the relative orientation of the input genome assembly along the pangenome graph.

**Figure 2.**
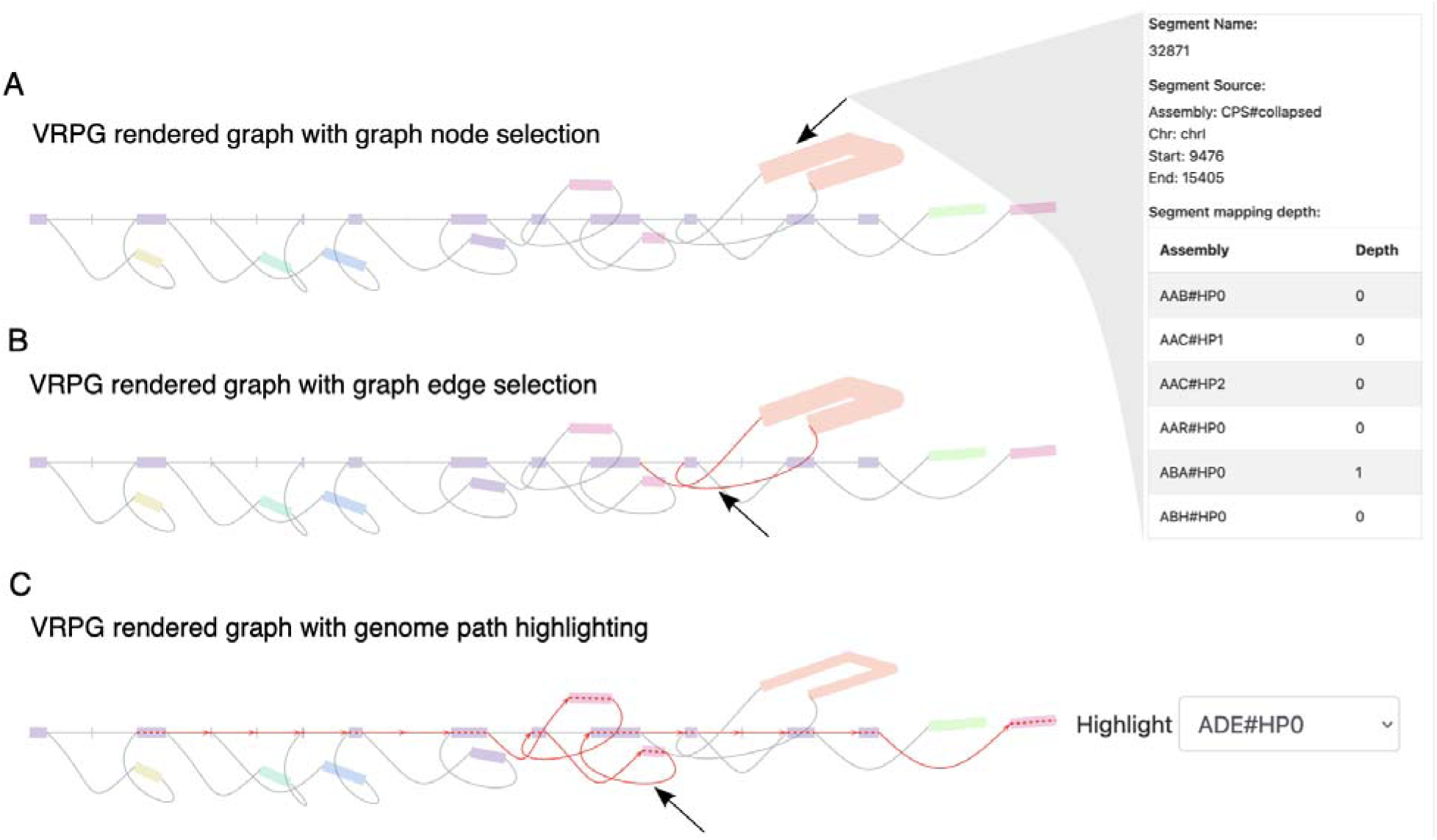
The reference pangenome graph visualization with VRPG. (A) VRPG visualization with the node s32871 selected by clicking on this graph node. (B) VRPG visualization with the two edges surround the node s32871 selected by clicking on these two graph edges. (C) VRPG visualization with the assembly-to-graph mapping path of the assembly ADE#HP0 highlighted by selecting ADE#HPD in the “Highlight” selection field.

### Node information inquiry

In VRPG, users can easily obtain the detailed genomic segment information of any node together with the node’s covered gene list and copy number status across all input assemblies by a simple click. Here the source genome assembly information specifies the genome assembly origin and coordinates (1-based) of the genomic segment that defines the corresponding node when building the pangenome graph. When preprocessing the input pangenome graph, VRPG will summarize the node-specific mapping depth by aligning each input genome assembly to the pangenome graph and calculate the assembly-to-graph mapping depth for all nodes encompassed in the graph. During visualization, the mapping depth of the query node will be reported for all input assemblies, where the mapping depth value reflects the actual copy number of the corresponding genomic segment that defines this node.

### Genomic variant representation in VRPG

One major motivation of developing a pangenome graph visualization tool is to provide an intuitive way to associate specific genomic variants to their graph representation. With VRPG and its reference-based projection and assembly path highlighting feature, this is very straightforward. In Figure 3, we exemplified how different types of genomic variants from a query genome relative to the reference genome were depicted by VRPG in the context of pangenome graph (Figure 3). As VRPG sequentially depicts nodes representing the reference genome according to their actual genomic coordinates along the center line, any highlighted query genome path that deviates this center line implies the occurrence of one or more genomic variants. The direction of the path reflects the connecting order from one node to another, which is especially useful when identifying directional variants such as inversions. The path will also be thickened if the highlighted genome traverses the path more than once, which implies the occurrence of duplications.

**Figure 3.**
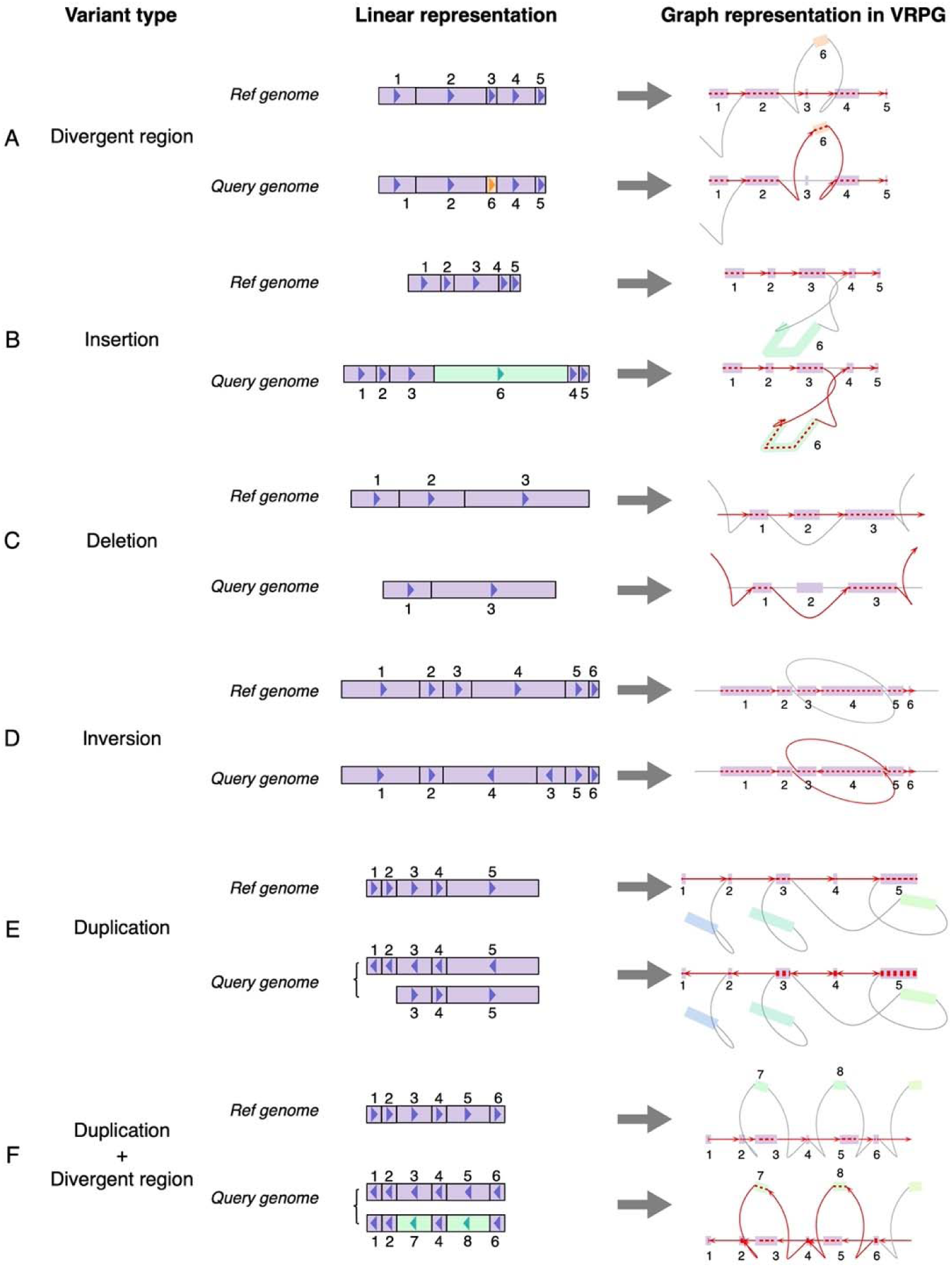
VRPG visualization for different types of genomic variants. For different types of genomic variants such as (A) Divergent region, (B) Insertion, (C) Deletion, (D) Inversion, (E) Duplication, and (F) Duplication with divergent region, their graph representations in VRPG are depicted with path highlighting for the reference and query genome respectively.

Application demonstration 1: visualizing pangenome graph derived from 163 yeast genome assemblies.

To demonstrate the application of VRPG in real world examples, we set up a dedicated webserver (https://www.evomicslab.org/app/vrpg/) to visualize the reference pangenome graph derived from 163 budding yeast genome assemblies. Here we employed Minigraph to construct a pangenome graph using the *Saccharomyces cerevisiae* reference assembly panel (ScRAP) that we recently assembled [16]. This yeast pangenome graph consists of 37,062 nodes and 52,756 edges, with a total length of 2,7190,479 bp.

### The flip/flop inversion

Based on this yeast pangenome graph, we used VRPG to visualize a famous flip/flop inversion region on the chromosome XIV (chrXIV) of the *S. cerevisiae* genome. This flip/flop inversion region is flanked by two 4.2-kb inverted repeat (IR) regions and remains polymorphic in *S*. *cerevisiae* and its sister species in the genus *Saccharomyces* [17]. For example, within *S. cerevisiae*, our previous study revealed that the Sake strain Y12 shared conserved synteny with the *S*. *cerevisiae* reference strain S288C within this region whereas the North American strain YPS128 showed an inverted form relative to S288C within this region instead [1] (Figure 4A). In accordance with this prior knowledge, by enabling the genome path highlight feature, we can recapture such polymorphic inversion with VRPG in the pangenome graph. As illustrated by VRPG, both S288C and Y12 revealed a simple linear assembly-to-graph path through the nodes along the central line, suggesting their reference-like sequence structure in this region (Figure 4B and 4C). In contrast, YPS128 showed an S-shape assembly-to-graph path in the flip/flop region, which implied the existence of an inverted sequence structure (Figure 4D).

**Figure 4.**
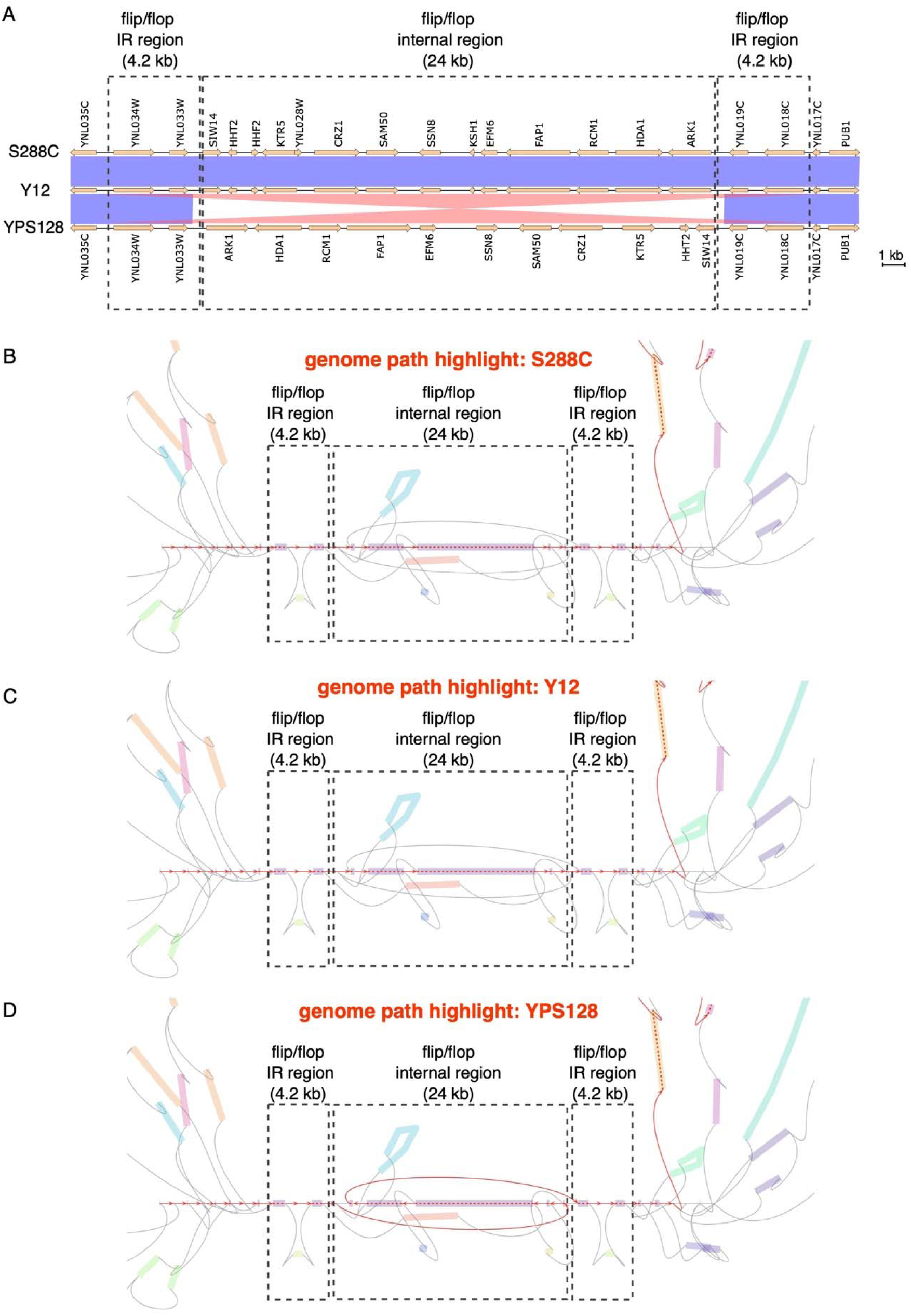
VRPG visualization for the chromosome XIV (chrXIV) flip/flop region in the yeast pangenome graph. (A) The genome sequence and synteny comparison among S288C, Y12, and YPS128 for the chrXIV flip/flop region, with the red and blue shades representing homologous regions shared with >97% sequence similarity (blue: same direction; red: reverse direction). (B-D) VRPG visualization for the yeast pangenome graph correspond to the query reference coordinate of chrXIV:569857-602365 with the genome path of S288C (B), Y12 (C), and YPS128 (D) highlighted respectively.

### The *DOG2* deletion

The yeast paralog gene pairs *DOG1* and *DOG2* encode 2-deoxyglucose-6-phosphate phosphatase involved in glucose metabolism. They are homologous to the human *PUDP* (*HDHD1*) gene, an anti-cancer treatment target for intervene the glycolytic metabolism of tumor cells [18]. Previously, we have identified a polymorphic deletion for the *DOG2* gene in the comparison of seven representative yeast strains using their T2T genome assemblies [1]. For example, this gene is present in strains like S288C, DBVPG6765, but absent in other strains such as DBVPG6044, SK1, and Y12 (Figure 5A). Here we used VRPG to visualize this gene presence/absence variation in the context of yeast pangenome graph (Figure 5B). With the strain S288C as the reference, we observed simple linear assembly-to-graph paths for S288C and DBVPG6765, suggesting their reference-like sequence structure in this region. Conversely, the assembly-to-graph paths for DPVPG6044, SK1 and Y12 clearly deviated from the reference path by bypassing the node representing the *DOG2* gene region.

**Figure 5.**
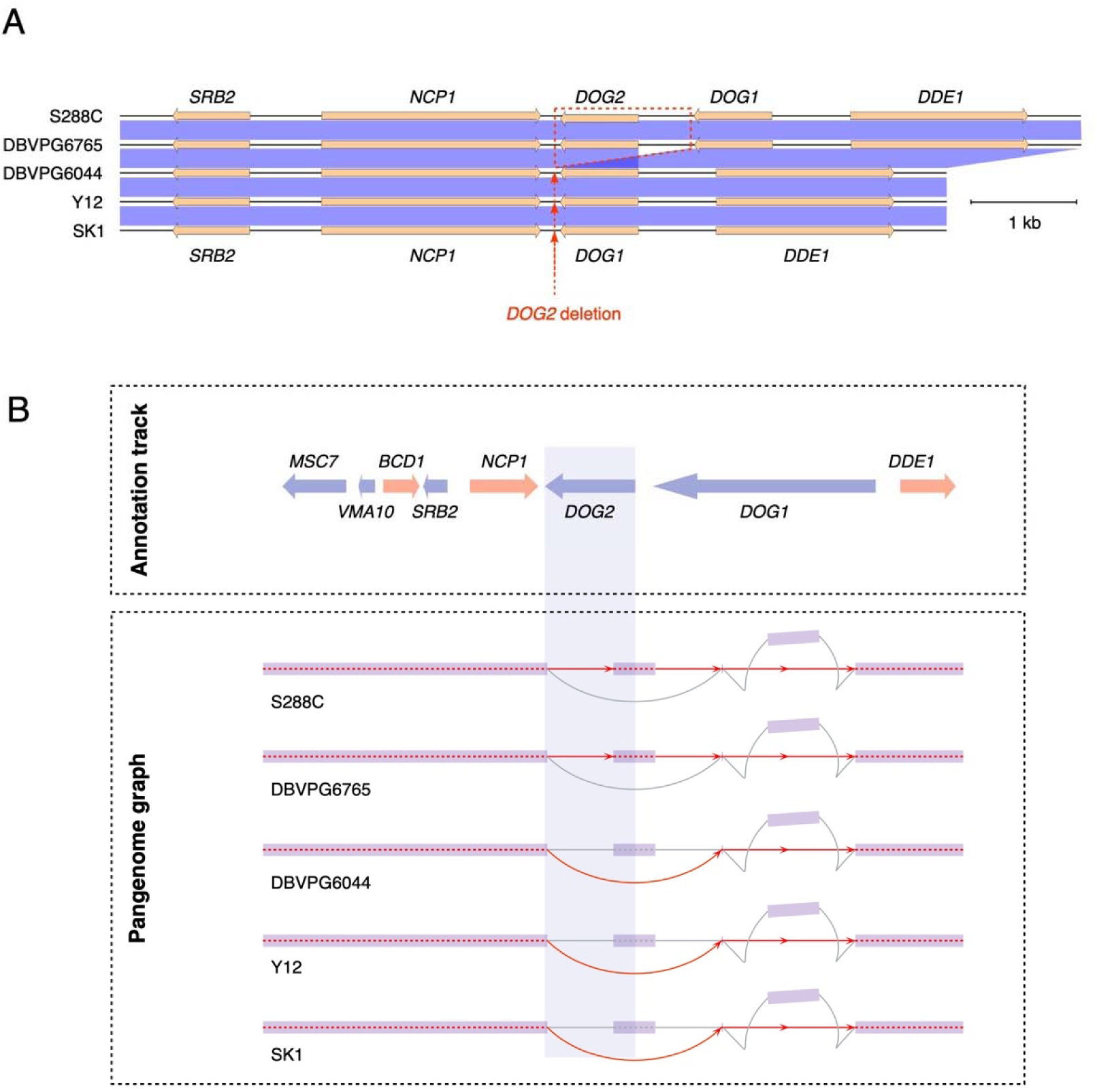
VRPG visualization for the polymorphic *DOG2* deletion in the yeast pangenome graph. (A) The genome sequence and synteny comparison among S288C, DBVPG6765, DBVPG6044, SK1 and Y12 for the *DOG2* region, with the blue shades representing homologous region shared with >97% sequence similarity. (B) VRPG visualization for the yeast pangenome graph correspond to the query reference coordinate of chrVIII:189131-197233 with the genome path of S288C, DBVPG6765, Y12, and SK1 highlighted respectively. The corresponding reference-based gene annotation track is displayed above.

### Application demonstration 2: visualizing pangenome graphs derived from 90 human genome assemblies

As a further proof for VRPG’s versatility and scalability, we retrieved the human pangenome graphs generated by the Human Pangenome Reference Consortium (HPRC) based on 90 human genome assemblies [19] and visualized them via our VRPG demonstration webserver (https://www.evomicslab.org/app/vrpg/). Starting with the same input genome set, HPRC built three human pangenome graphs with Minigraph, Minigraph-CACTUS, and PGGB respectively. The HPRC human pangenome graphs built by Minigraph consists of 391,950 nodes and 566,204 edges, with a total length of 3,198,196,033 bp. In comparison, the total node and edge numbers of graphs built by Minigraph-CACTUS (81,415,956 nodes and 112,955,105 edges) and PGGB (110,884,673 nodes and 154,756,169 edges) are substantially larger as small variants such as SNV and INDELs are fully considered by Minigraph-CACTUS and PGGB during their graph construction whereas Minigraph predominantly considers larger variants such as SVs.

### The *DSCAM* intronic inversion

For the demonstration, we used VRPG to visualize an inversion locate in the intron 32 of the *DSCAM* gene[20] (Figure 6). This inversion is mediated by two inverted 6.0-kb L1 elements, forming a structure much like the yeast chrXIV flip/flop inversion described above (Figure 6A). *DSCAM* gene belongs to the immunoglobulin superfamily of cell adhesion molecules (Ig-CAMs) and functions in central and peripheral nervous system development. *DSCAM* gene is a clinically relevant gene with significant association with Down syndrome (DS) and congenital heart disease (DSCHD) [21,22]. This *DSCAM* inversion is segregate between the current human reference genome GRCh38 and the telomere-to-telomere (T2T) assembly CHM13 (Figure 6B). When highlighting the path of the CHM13 genome assembly, this inversion is clearly visible in the pangenome graph built by Minigraph (Figure 6C). For the pangenome graph built by Minigraph-CACTUS and PGGB, the graph layouts are much more complex due to the incorporation of small variants (i.e., variants <50bp) by Minigraph-CACTUS and PGGB. This is where the layout simplification function of VRPG can help, which serves to generate a much simpler graph layout by hiding small variant related nodes. We found the *DSCAM* inversion in CHM13 popped up clearly with a characteristic S-shape path in the simplified Minigraph-CACTUS pangenome graph (Figure 6D). In the PGGB graph, the pattern is more complex even with layout simplification, showing loop-like structure in the inversion region (Figure 6E), which highlights the methodological difference between PGGB and the other two graph builders (i.e., Minigraph and Minigraph-CACTUS) during the graph building process.

**Figure 6.**
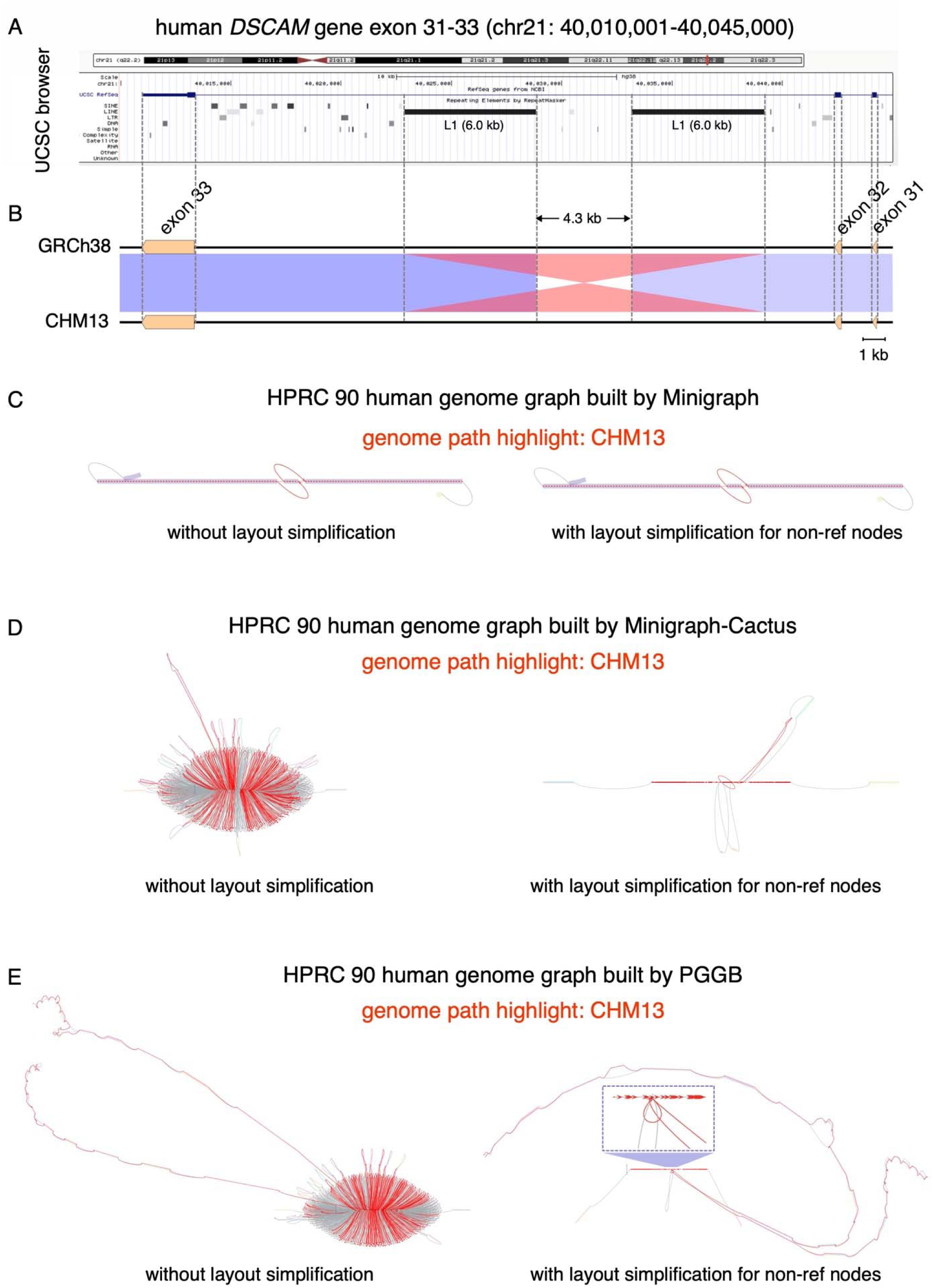
VRPG application demonstration with the human pangenome graphs for the *DSCAM* intronic inversion. (A) The UCSC genome browser view for the *DSCAM* exon 31-33 region at chr21:40,010,001-40,045,000 with annotation tracks for chromosome ideogram, gene structure, repetitive sequence features. (B) The genome sequence and synteny comparison between the human reference genome GRCh38 and the telomere-to-telomere (T2T) genome CHM13 for the *DSCAM* exon 31-33 region, with the red and blue shades representing homologous regions shared with >98% sequence similarity. (C-E) VRPG visualization for the *DSCAM* inversion in the human pangenome graphs derived from Minigraph (C), Minigraph-CACTUS (D), and PGGB (E), with the genome path of CHM13 highlighted respectively.

### The *CR1* intragenic deletion

The *CR1* gene (also known as *CD35*) is a complement activation receptor gene that is strongly associated with the Alzheimer disease[23,24]. This gene is structurally polymorphic in human populations, bearing an intragenic duplication region whose copy number correlates with the risk of Alzheimer disease[25]. An 18.6-kb intragenic deletion has been reported for *CR1* in a recent comparison between the human GRCh38 reference genome and the CHM13 T2T genome, with the latter one hosting the shorter version[26]. Here we took a closer examination for this deletion in the context of its nearby sequence homology and confirmed that this deletion locates within the previously reported intragenic duplication region (Figure 7A-B). Next, we used VRPG to visualize this *CR1* intragenic deletion based on the human pangenome graphs built by Minigraph, Minigraph-CACTUS, and PGGB respectively (7C-E). By comparing the highlighted paths of GRCh38 and CHM13 in VRPG, we can clearly see the signal of deletion in both Minigraph and Minigraph-CACTUS pangenome graphs. As for the pangenome graph built by PGGB, VRPG also depicted notable path difference between GRCh38 and CHM13 (note those extra red paths highlighted for CHM13), although the PGGB graph is topologically too complex to reflect the deletion in an intuitive way.

**Figure 7.**
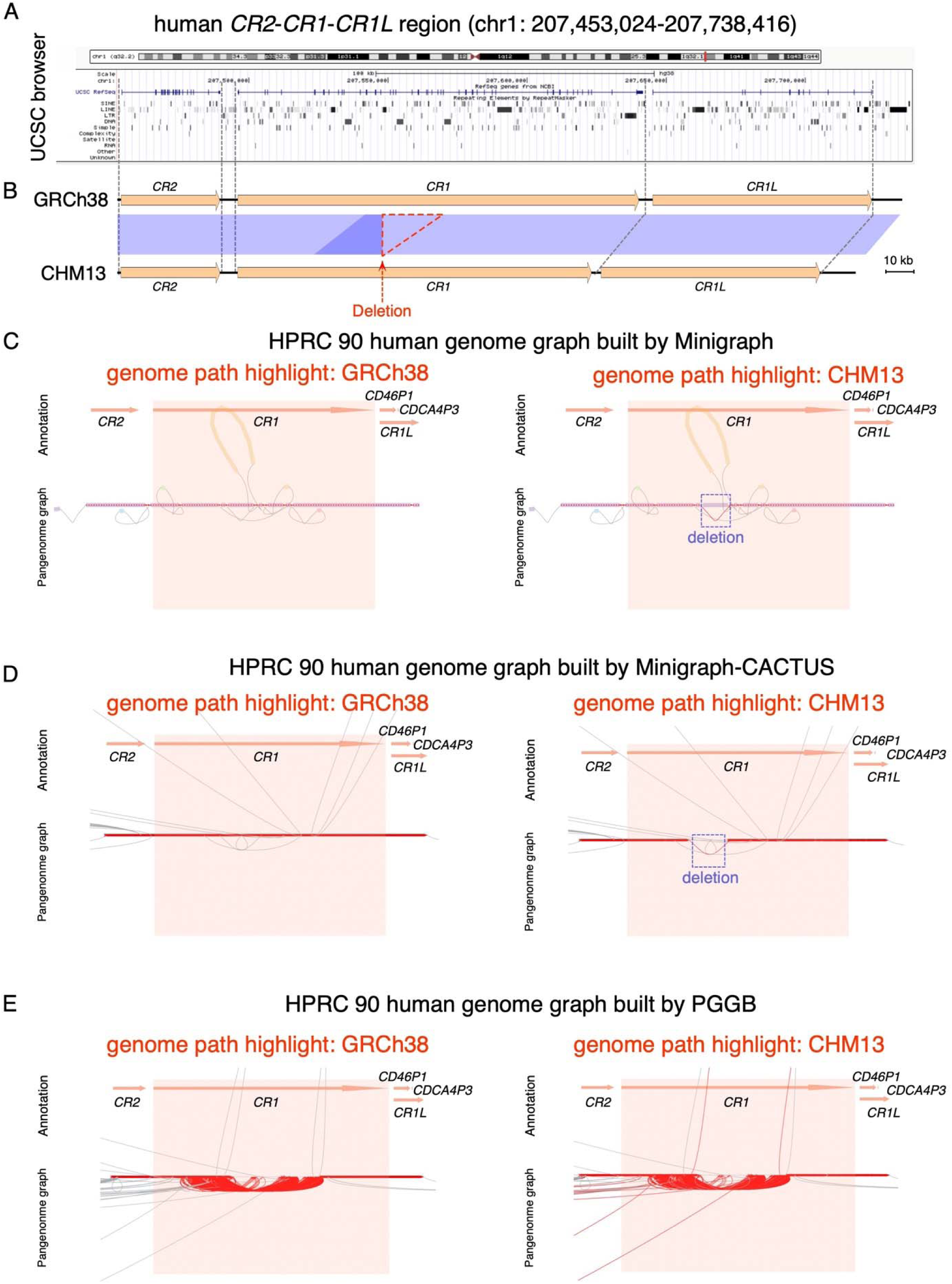
VRPG application demonstration with the human pangenome graphs for the *CR1* intragenic deletion. (A) The UCSC genome browser view for the *CR2*-*CR1-CR1L* region in chr1:207,453,024-207,738,416 (GRCh38 reference genome) with annotation tracks for chromosome ideogram, gene structure, repetitive sequence features. (B) The genome sequence and synteny comparison between the human reference genome GRCh38 and the T2T genome CHM13 for the *CR2*-*CR1-CR1L* region, with the blue shades representing homologous regions shared with >98% sequence similarity (blue: same direction; red: reverse direction). (C-E) VRPG visualization for the *DSCAM* inversion in the human pangenome graphs derived from Minigraph (C), Minigraph-CACTUS (D), and PGGB (E), with the genome paths of GRCh38 and CHM13 highlighted respectively. Non-ref node simplification was triggered when visualizing the Minigraph-CACTUS and PGGB graph.

### Limitations

Although efforts have been taken, there are still several limitations for VRPG at the current stage. For example, although VRPG generally keep the reference nodes well-scaled to their actual genomic sizes, the rendered sizes of non-reference peripheral nodes could be substantially altered for displaying convenience. Besides, only annotation features specified in GFF3 format is currently supported, whereas the supports for other widely used genomic formats such as BED, VCF, and WIG are yet to be added. Improvements that tackle these limitations are planned to be implemented in near future, with the help from the ever-growing pangenome graph community.

## Discussion

As genome sequencing technologies keep progressing towards longer read length and higher per-base accuracy, generating reference-quality genomes at a population scale is becoming increasingly affordable [27]. Along with this tread, researchers began to develop new computational frameworks to accommodate and analyze such growing collection of high-quality genomes with better efficiency. Pangenome graph serves as a compact yet extendable data structure that describes population-level genetic diversity at both base and structure level via a graph representation [4]. This is a highly active research field with new tools and applications rolling out very quickly. One can envision that many genomics analyses that currently relies on conventional linear reference genome will eventual be performed based on pangenome graphs in near future. A better understanding of how different layouts and topologies of pangenome graphs correspond to the actual genomic variation in the population is of critical for correctly interrogating and interpretating pangenome-graph-based analysis results. In this study, we developed VRPG, a web-based interactive pangenome graph visualization framework with full scalability, capable of rendering complex pangenome graphs derived from hundreds of genome assemblies. In addition to intuitive graphical visualization, VRPG also provides biologically relevant features that directly reflects the sequence and copy number variation of the genome assemblies that comprise the graph. As demonstrated with real world examples based on yeast and human pangenome graphs, we expect VRPG to become a highly useful visualization tool to help researchers to explore the full power of pangenome graph.

## Conclusions

Here we present VRPG, a web-based pangenome graph visualization framework. In addition to enabling interactive visualizing and analyzing pangenome graph on the fly, VRPG also shines in providing an intuitive interface that connects linear genome assemblies and their embedded genomic variants to the pangenome graph.

## Availability

VRPG is free for use under the MIT license, with the source code available at https://github.com/codeatcg/VRPG. The real-case demonstration of VRPG on yeast and human pangenome graph is hosted at https://www.evomicslab.org/app/vrpg/.

## Materials and Methods

### Naming scheme of input assemblies

The naming scheme of input assemblies in VRPG follows a relaxed version of the PanSN prefix naming specification (https://github.com/pangenome/PanSN-spec). The input genome assembly identifier consists of three parts: sample tag, delimiter, and haplotype tag. The sample tag and haplotype tag can be any strings consist of letters and/or numbers, while the delimiter can be any character (e.g., “#” or “:”).

### Input pangenome graph specification

In a pangenome graph, a node represents a directed genomic segment while an edge represents the relative order and direction of the two inter-connected nodes. VRPG natively works with pangenome graph files in the rGFA format, which can be straightforwardly constructed using Minigraph[6]. Compared with pangenome graph encoded in other formats, the rGFA-formatted pangenome graph inherently tracks the origin of each node in terms of its location and direction from the input assembly, which comes handy when analyzing and interpretating the corresponding pangenome graph. In addition, VRPG offers further support for genome graphs encoded in the GFAv1 format with a pre-shipped sub-module named gfa2view, which can transform the GFAv1 format into an rGFA-like format that encodes segment coordinate information as well.

### GFAv1 to rGFA-like transformation

GFAv1 is a widely used pangenome graph format. Unlike the rGFA-formatted pangenome graph, there is no pre-defined reference genome for establishing the coordinate system when constructing the pangenome graph in the GFAv1 format. Therefore, to accommodate GFAv1-formated graph with VRPG, a user defined reference genome is required. When there are multiple copies of highly similar segments (e.g., in the case of segmental or tandem duplication) in a given genome assembly, some pangenome graph builders such as PGGB tends to collapse them into a single node, which is more efficient in compressing genomic information but also breaks the linearity of the reference genome coordinate system. To walkaround this challenge, the gfa2view module of VRPG will traverse the reference path to find the collapsed nodes, which are nodes been traversed for two or more times and insert the corresponding “shadow nodes” and their associated edges into the graph to restore the original reference coordinates for the given node. In this way, the reference genome rendered by VRPG by sequentially chaining through reference nodes still keeps linearity and the genomic regions collapsed by the original pangenome graph builder (e.g., PGGB) can still be traced with VRPG’s path highlighting function. The resulting graph with these added shadow nodes is recorded in a rGFA-like format, which can be used for VRPG as the input file for visualization.

### Node and edge rendering

Representative genomic segments are denoted by graph nodes and further illustrated as colored blocks in the view window. For nodes representing the reference genome (pre-defined when building the reference pangenome graph), their block sizes are generally proportional to the actual size of the corresponding genomic segments. For nodes representing non-reference assemblies, their corresponding blocks can only roughly reflect the actual segment sizes. If the segment size (denoted as “S” herein) is smaller than 2 kb, the node will be illustrated as a rectangular block. If the segment size is greater than 2 kb, the node will be represented by a curved block where the corner count of the curved block is proportional to the multiples of 2-kb determined by S. The connections between different genomic segments are represented by graph edges and further illustrated as directed lines. For the cleanness and efficacy of rendering, additional graph simplification algorithms are further applied to trim off those peripheral nodes that are far away from the reference nodes (e.g., when traversing depths >= 10) as well as optionally to hide small nodes (e.g., those representing genomic variants that are < 50 bp) in the graph. These features are especially helpful when visualizing genome graphs built by Minigraph-CACTUS and PGGB where all base-level variants such as SNV and INDEL are coded as nodes.

### Efficient access to the graph

VRPG implemented a block index system to enable quick access to the subgraph corresponding to any given region along the pre-defined reference genome. The reference genome was first subdivided into blocks with each block containing 2000 reference nodes. For each block, a breadth-first search algorithm was implemented to find all edges and non-reference nodes associated with the reference nodes within the corresponding block and a block index corresponding to these edges and nodes was created. To reduce the search space and accelerate the indexing process, VRPG only uses edges that have no joins with all other chromosomes/scaffolds/contigs during indexing. Based on the indexed nodes and edges, an ordered block index array for each assembly path was created. Based on such block index system, VRPG can efficiently locate the subgraph associated with the query region for rendering.

### Yeast reference pangenome graph construction

The yeast reference pangenome graph was constructed by this study. This graph was built upon 163 yeast genome assemblies from 142 strains, with some heterozygous/polyploid strains having both phased and collapsed assemblies. Briefly, we took the *Saccharomyces cerevisiae* reference genome (version: R64-1-1; denoted as SGDref) retrieved from *Saccharomyces* genome database (SGD; https://www.yeastgenome.org/) as well as 162 assemblies from our recently released *Saccharomyces cerevisiae* Reference Assembly Panel (ScRAP) [16] to construct reference pangenome graph. Minigraph [6] was used for this graph construction with the command ‘minigraph -cxggs -l 5000’. With the SGDref as the reference genome, we incrementally added those 162 ScRAP assemblies into the graph according to their phylogenetic distances to SGDref. The phylogenetic distances employed here were extracted from the phylogenetic tree of these 163 input genomes built upon their concatenated 1-to-1 orthologous gene matrix (See [16] for details). Regarding the haplotype tag, for the yeast reference pangenome graph, we used “HP0” to denote haplotypes of haploid or homozygous diploid strains, while using “collapsed”, “HP1”, and “HP2” to denote collapsed, or the two phased haplotypes of heterozygous diploid strains.

### Human reference pangenome graph construction

Human Pangenome Reference Consortium (HPRC) constructed three human pangenome graphs using Minigraph, Minigraph-CACTUS, and PGGB respectively based on 90 genome assemblies from 46 samples [19]. Aside from the classic (GRCh38) and T2T (CHM13) human reference genome assemblies, the other 44 samples all have two fully phased genome assemblies. We retrieved the human Minigraph-CACTUS and PGGB pangenome graph from the HRPC’s Github page (https://github.com/human-pangenomics/hpp_pangenome_resources) and downloaded a slightly updated version of the Minigraph pangenome graph provided by Heng Li (https://zenodo.org/record/6983934#.Y767A3ZByUk)[28]. For these human reference pangenome graphs, “0” was used to denote haplotypes of haploid or collapsed samples while “mat” and “pat” were used to denote the maternal and paternal haplotypes of the phased diploid samples.

### Software implementation and web deployment

The backend of VRPG is implemented in C++ and Python3 based on the Django framework, while both D3.js (https://github.com/d3/d3) and WebCola (https://github.com/tgdwyer/WebCola) were employed at the frontend for data visualization. Bootstrap and jQuery were further used for interactive query and rendering. The demonstration webserver is deployed via an Alibaba Simple Application Server with 2 CPU, 4 Gb RAM, and 80 Gb ESSD data storage.

### Linear genome comparison and visualization for the demonstrated examples

The genomic regions of the demonstrated loci (the chrXIV flip/flop inversion and *DOG2* for yeast as well *DSCAM* and *CR1* for human) linear genome were extracted from the corresponding genome assemblies and subsequently compared and plotted with EasyFig[29].

## Acknowledgements

We are grateful to Dr. Jing Li for valuable feedback on the manuscript.

## Author contributions

J.-X. Y. conceived the study. M.Z. wrote the software and analyzed the results. J.-X. Y. and M.Z. wrote the paper. All authors reviewed and contributed to the final version of the paper. Correspondence to J.-X. Y.

